# Development of microsatellite markers for a giant water bug, *Appasus japonicus*, distributed in East Asia

**DOI:** 10.1101/2020.05.25.115501

**Authors:** Tomoya Suzuki, Akira S. Hirao, Masaki Takenaka, Koki Yano, Koji Tojo

**Affiliations:** Faculty of Science, Shinshu University, Asahi 3-1-1, Matsumoto, Nagano 390-8621, Japan; Sugadaira Research Station, Mountain Science Center, University of Tsukuba, Sugadairakogen 1278-294, Ueda, Nagano 386-2204, Japan; Faculty of Symbiotic Systems Science, Fukushima University, Kanayagawa 1, Fukushima, Fukushima 960-1296, Japan; Division of Evolutionary Developmental Biology, National Institute for Basic Biology, Nishigonaka 38, Myodaiji, Okazaki, Aichi 44-8585, Japan; Department of Mountain and Environmental Science, Interdisciplinary Graduate School of Science and Technology, Shinshu University, Asahi 3-1-1, Matsumoto, Nagano 390-8621, Japan; Institute of Mountain Science, Shinshu University, Asahi 3-1-1, Matsumoto, Nagano 390-8621, Japan

**Author notes:** Corresponding authors Koji Tojo, Tomoya Suzuki.

**Keywords:** endangered species, giant water bug, genetic variation, SSR, Ion PGM

## Abstract

We developed microsatellite markers for *Appasus japonicus* (Hemiptera: Belostomatidae). This belostomatid bug is distributed in East Asia (Japanese Archipelago, Korean Peninsula, and Mainland China), and often listed as endangered species in the ‘Red List’ or the ‘Red Data Book’ at the national and local level in Japan. Here we describe twenty novel polymorphic microsatellite loci developed for *A. japonicus*, and marker suitability was evaluated on 56 individuals from four *A. japonicus* populations (Nagano, Hiroshima, and Yamaguchi prefecture, Japan, and Chungcheongnam-do, Korea). The number of alleles per locus ranged 1–12 (mean = 2.5), and average observed and expected heterozygosity, and fixation index per locus were 0.270, 0.323, and 0.153, respectively. The 20 markers described here will be useful for investigating the genetic structure of *A. japonicus* populations, which can contribute in population genetics studies of this species.

---

Freshwater biodiversity, including that of aquatic invertebrates, is the overriding conservation priority of the International ‘Water for Life’ Decade for Action (Dudgeon et al., 2006; Doi et al., 2017). *Appasus japonicus* is an aquatic insect, which is distributed throughout the Japanese Archipelago, Korean Peninsula, and Mainland China. This species is often listed as endangered species in the ‘Red List’ or the ‘Red Data Book’ at the national and local level (Ministry of the Environment, Japan, 2006). Their evolutionary history is revealed by our previous study using mtDNA COI and 16S rRNA regions, and three largely divided genetic lineages were identified within this species (Suzuki et al., 2013, 2014). Furthermore, “back dispersal” of *A. japonicus*, i.e., dispersal from the Japanese Archipelago to Eurasian continent, was suggested from our previous study (Suzuki et al., 2014). However, more fine-scale analyses, like a population genetic analysis, have not been conducted. The microsatellite marker is one of the most useful tools for identifying the population genetic structure and many studies using microsatellite markers for the fine-scale population genetic analyses (e.g., Phillipsen and Lytle, 2013; Phillipsen et al., 2015; Hirao et al., 2017; Komaki et al., 2017). Furthermore, the information of the population genetic structure is very important for conservation of organisms. Therefore, in this study, we developed twenty microsatellite markers for *A. japonicus*, and evaluated marker suitability using for population genetic analyses.

Microsatellite markers were developed for *A. japonicus* using the Ion PGM system (Life Technologies). Library preparation and PGM sequencing were conducted the Sugadaira Montane Research Station, Mountain Science Center, University of Tsukuba, Japan. Total genomic DNA was extracted from the ethanol-preserved tissue of specimens which collected in Matsumoto, Nagano, and purified using the DNeasy Blood & Tissue Kit (QIAGEN, Hilden) according to the manufacturer’s instructions. The concentration of genomic DNA was quantified by a Qubit 2.0 Fluorometer (Life Technologies), and 13.6 ng/μL of DNA was used for the following processes. The genomic DNA was sheared to approximately 350–450 bp by Ion Shear Plus Reagents (Life Technologies), and the adapter ligation, nick-repair, and purification of the ligated DNA was conducted using an Ion Plus Fragment Library Kit (Life Technologies). After size selection (target insert sizes 300–400 bp) was performed by an E-Gel Agarose Gel Electrophoresis System (Life Technologies), library amplification was conducted using an Ion Plus Fragment Library Kit (Life Technologies). The library was assessed and quantified using a Bioanalyzer (Agilent Technologies, Palo Alto, California, USA), and then diluted to 8 pM for template preparation using an Ion PGM Template OT2 400 kit (Life Technologies) and enriched. Sequencing was performed by an Ion PGM Sequencing 400 kit (Life Technologies) using 850 flows on the Ion 314 Chip V2 (Life Technologies) according to the manufacturer’s protocol. After sequencing, single processing and base-calling were performed using TorrentSuite 3.6 (Life Technologies), and a library-specific FASTQ file was generated. The data sets were collated and applied to the QDD bioinformatics pipeline (Meglécz et al., 2010) to filter sequences containing microsatellites with appropriate flanking sequences to define PCR primers. QDD detected 10,760 loci, each containing a microsatellite consisting of at least five repeats. A total of 50 primer pairs were obtained for screening. Twenty primer pairs showing clear peak patterns were selected after an initial primer screening using 8 samples from Matsumoto, Nagano population, and 8 samples from Shimonoseki, Hiroshima population (Table 1).

**Table 1.**
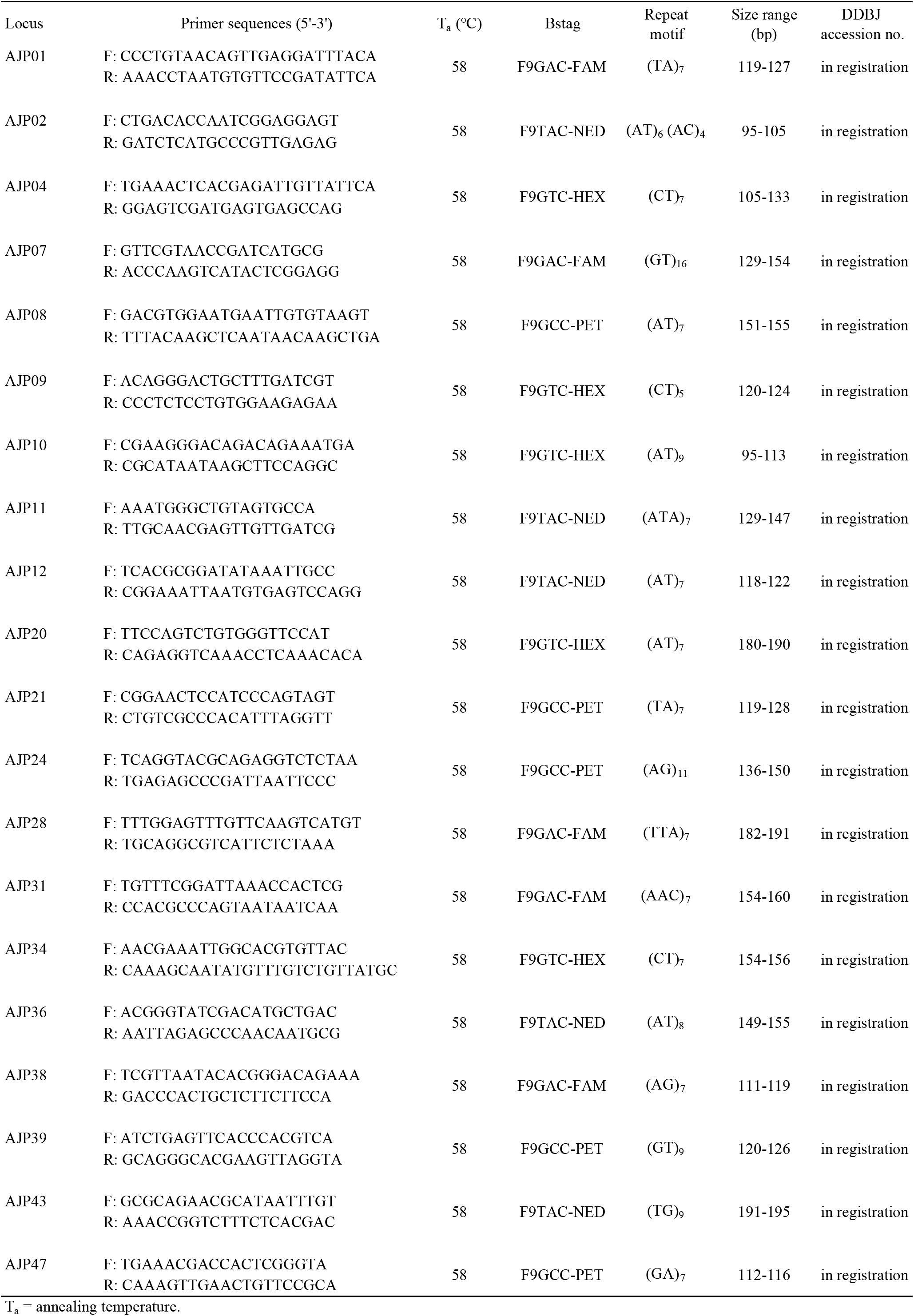
Characteristics of 20 microsatellite primers developed for *Appasus japonicus*.

To test the genetic variation of the 20 selected microsatellite loci, 20 samples from Matsumoto, Nagano, 10 samples from Mihara, Hiroshima, and 10 samples from Shimonoseki, Yamaguchi were used. PCR amplification with fluorescent dye-labeled primers was performed using a protocol described by Shimizu and Yano (2011). PCR amplification was done in 10 μL reactions using the KOD FX Neo DNA polymerase (TOYOBO, Osaka, Japan). Each reaction contained the following components: 1 μL of total genomic DNA, 4.8 μL of 2 × buffer, 1.6 μL of 2.0 mM dNTP mix, 0.05 μL of forward primer, 0.2 μL of reverse primer, 0.05 μL of fluorescent dye-labeled primer and 2.3 μL of SQ. The PCR protocol was: 94°C for 2 min; 30× (98°C for 10 sec, 58°C for 10 sec, and 68°C for 30 sec); 68°C for 5 min. We labeled BStag primers with the following fluorescent dyes: F9GAC-FAM (5’-CTAGTATCAGGACGAC-3’), F9GTC-HEX (5’-CTAGTATGAGGACGTC-3’), F9TAC-NED (5’-CTAGTATCAGGACTAC-3’), F9GCC-PET (5’-CTAGTATTAGGACGCC-3’), and F9CCG-FAM (5’-CTAGTATTAGGACCCG-3’). Product sizes were determined using an ABI 3130xl Genetic Analyzer and GeneMapper software (Applied Biosystems) with GeneScan 500 LIZ dye Size Standard v2.0 (Applied Biosystems). We calculated observed heterozygosity (*H*_O_), expected heterozygosity (*H*_E_) and inbreeding coefficients (*F*_IS_) using GenAlEx 6.5 (Peakall and Smouse, 2012). We also tested deviation from Hardy–Weinberg equilibrium and linkage disequilibrium among the polymorphic loci using GENEPOP 4.7 (Rousset, 2008).

As a result, all 20 microsatellite markers, which were developed in this study had meaningful polymorphism. 17 loci were stably amplified and genotyped in Nagano population, 15 loci were stably amplified and genotyped in Hiroshima and Yamaguchi population, and 11 loci were stably amplified and genotyped in Chungcheongnam-do population (Table 2). The number of alleles across per locus the four populations was 1–12 (mean = 2.5). Four and three loci were not polymorphic in the Hiroshima and Yamaguchi population, respectively (Table 2). The ranges of *H*_O_, *H*_E_ and *F*_IS_ per locus were 0.000–0.800 (mean = 0.270), 0.000–0.900 (mean = 0.323), and −0.414–1.000 (mean = 0.153), respectively (Table 2).

**Table 2.**
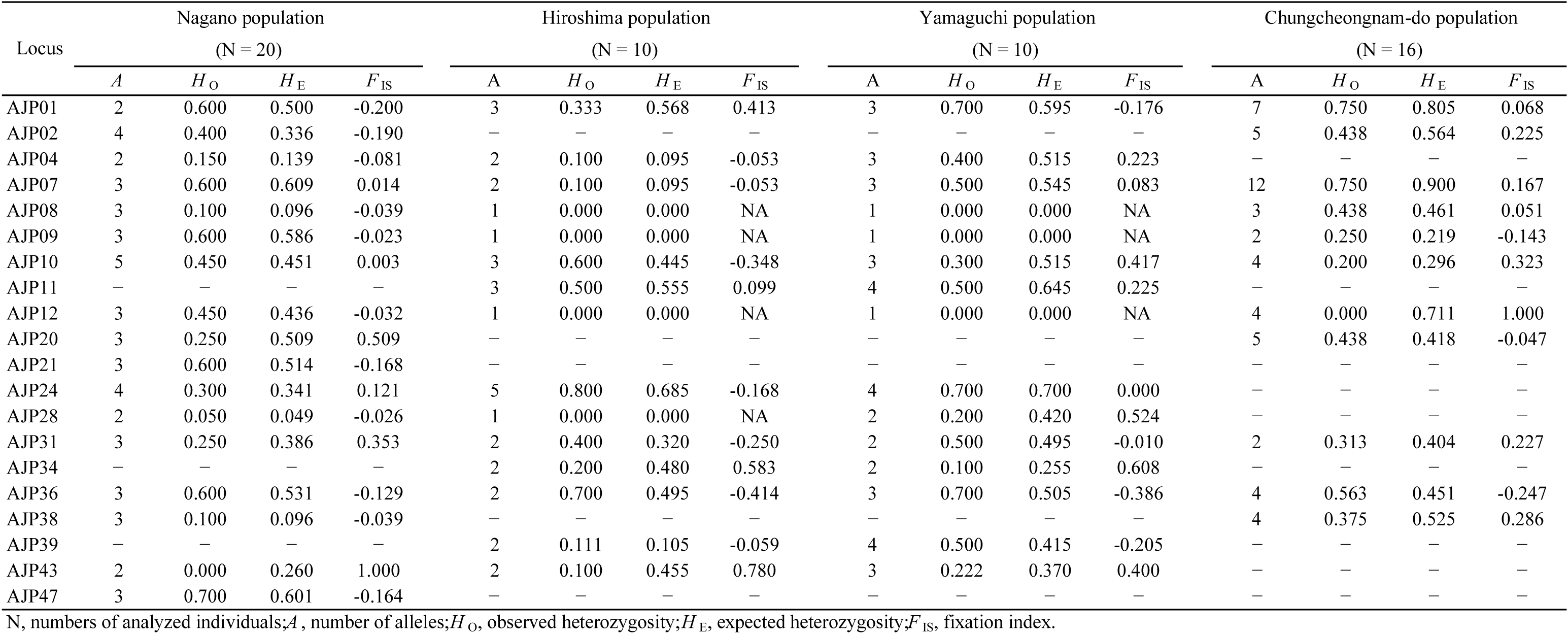
Genetic variation of the 20 micrisatellite loci for two populations of *Afppasus japonicus*.

In conclusion, we sequenced *A. japonicus* using Ion PGM and found microsatellite regions. Based on these data, we developed 20 polymorphic microsatellite markers for this species. These polymorphic markers are the first developed for *A. japonicus*. *A. japonicus* has a high potential as a model organism for the study of arthropod evolution (e.g. speciation, evolution of a paternal care system). These microsatellite markers are useful for elucidating broad- and fine-scale population genetic structure and evolution of unique paternal care mating systems in *A. japonicus*, not only conservation genetics research.

## Acknowledgements

We are sincerely thankful to Professor Emeritus K. Suzuki of Shinshu University for arranging the study environment. We also express our thanks to Dr. R. Saito (Fukushima Prefectural Centre for Environmental Creation), and Ms. K. Momose (Shinshu University) for their cooperation with the field research and collection of specimens, and Dr. H. Matsumura (Shinshu University) for providing facilities for genetic analysis. This study was supported by JSPS KAKENHI Grant Numbers JP20687005, JP23657046, JP16K14807 (K.T.), 285211031 (K.T.), and JP26891010, JP19K16209 (T.S.), the River Fund of The River Foundation, Japan [27-1215-013 (K.T.), and 27-1263-012 (T.S.)], and the Sasakawa Scientific Research Grant (26-529; T.S.).

